# The drivers of dark diversity in the Scandinavian tundra are metric-dependent

**DOI:** 10.1101/2023.02.17.528269

**Authors:** Lore Hostens, Koenraad Van Meerbeek, Dymphna Wiegmans, Keith Larson, Jonathan Lenoir, Jan Clavel, Ronja Wedegärtner, Amber Pirée, Ivan Nijs, Lembrechts Jonas J.

## Abstract

**Aim:** Dark diversity refers to the set of species that are not observed in an area but could potentially occur based on suitable local environmental conditions. In this paper, we applied both niche-based and co-occurrence-based methods to estimate the dark diversity of vascular plant species in the subarctic tundra. We then aimed to unravel the drivers explaining (1) why some locations were missing relatively more suitable species than others, and (2) why certain plant species were more often absent from suitable locations than others.

**Location:** The Scandinavian tundra around Abisko, northern Sweden.

**Methods:** We calculated the dark diversity in 107 plots spread out across four mountain trails using four different methods. Two niche-based (Beals’ index and hypergeometric method) and two co-occurrences-based (climatic niche model and climatic niche model followed by species-specific threshold) methods. This was then followed by multiple generalized linear mixed models and general linear models to determine which habitat characteristics and species traits contributed most to the dark diversity.

**Results:** The study showed a notable divergence in the predicted drivers of dark diversity depending on the method used. Nevertheless, we can conclude that plot-level dark diversity was generally 18% higher in areas at low elevations and 30% and 10% higher in areas with a low species richness or low levels of habitat disturbance, respectively.

**Conclusion:** Our findings call for caution when interpreting statistical findings of dark diversity estimates. Even so, all analyses point towards an important role for natural processes such as competitive dominance as main driver of the spatial patterns found in dark diversity in the northern Scandes.

## Introduction

Terrestrial ecosystems are increasingly affected by land-use and climate change, leading to large-scale biodiversity loss and community turnover (Theurillat & Guisan, 2001; Mooney et al., 2009; Newbold et al., 2015). Biodiversity plays an important role in ecosystem health and its loss alters ecosystem function (Hooper et al., 2012; Tilman et al., 2014). While most research has focused on the set of species that occur in an area, much less attention has gone to those species that are missing but could potentially inhabit the area (Pärtel et al., 2011). Nevertheless, to get a better understanding of community patterns and their underlying processes, such species absences hold viable additional information (Pärtel, 2014). Knowing which species from the regional species pool are absent within a given locality and identifying why, can help fine-tune conservation planning (Lewis et al., 2017). For example, if many of the absent – yet expected based on climate conditions – species are dispersal limited or cannot access the focal area due to strong dispersal barriers (i.e., habitat fragmentation), then some form of facilitated dispersal through assisted migration or actions to restore habitat connectivity is needed to restore biodiversity. However, if the nutrient conditions in the soil of the focal area are unsuitable for many of the missing species, then only providing assisted migration towards climatically suitable locations or restoring suitable climatic corridors would not be sufficient as restoration measures.

Species belonging to the missing part of the environmentally filtered regional species pool are defined as the so-called “dark diversity” (see Figure 1a), a term introduced by Pärtel et al. (2011). To be part of the dark diversity, the absent species must have a reasonable probability of dispersing to the area (i.e., by belonging to the regional species pool) and its ecological requirements (depending on the methodology either only its climatic or all environmental requirements) must match the local conditions (Pärtel, 2014). As a result, species that are present in the regional surroundings of the focal locality can be locally missing because they have a lower competitive ability, are dispersal limited, are ill-adapted to abiotic conditions, or due to stochastic processes (Riibak et al., 2015). Understanding how extrinsic abiotic conditions and intrinsic species characteristics related to competition and dispersal abilities influence a species’ absence can consequently give a better view of the community assembly (Belinchón et al., 2020).

**Figure 1:**
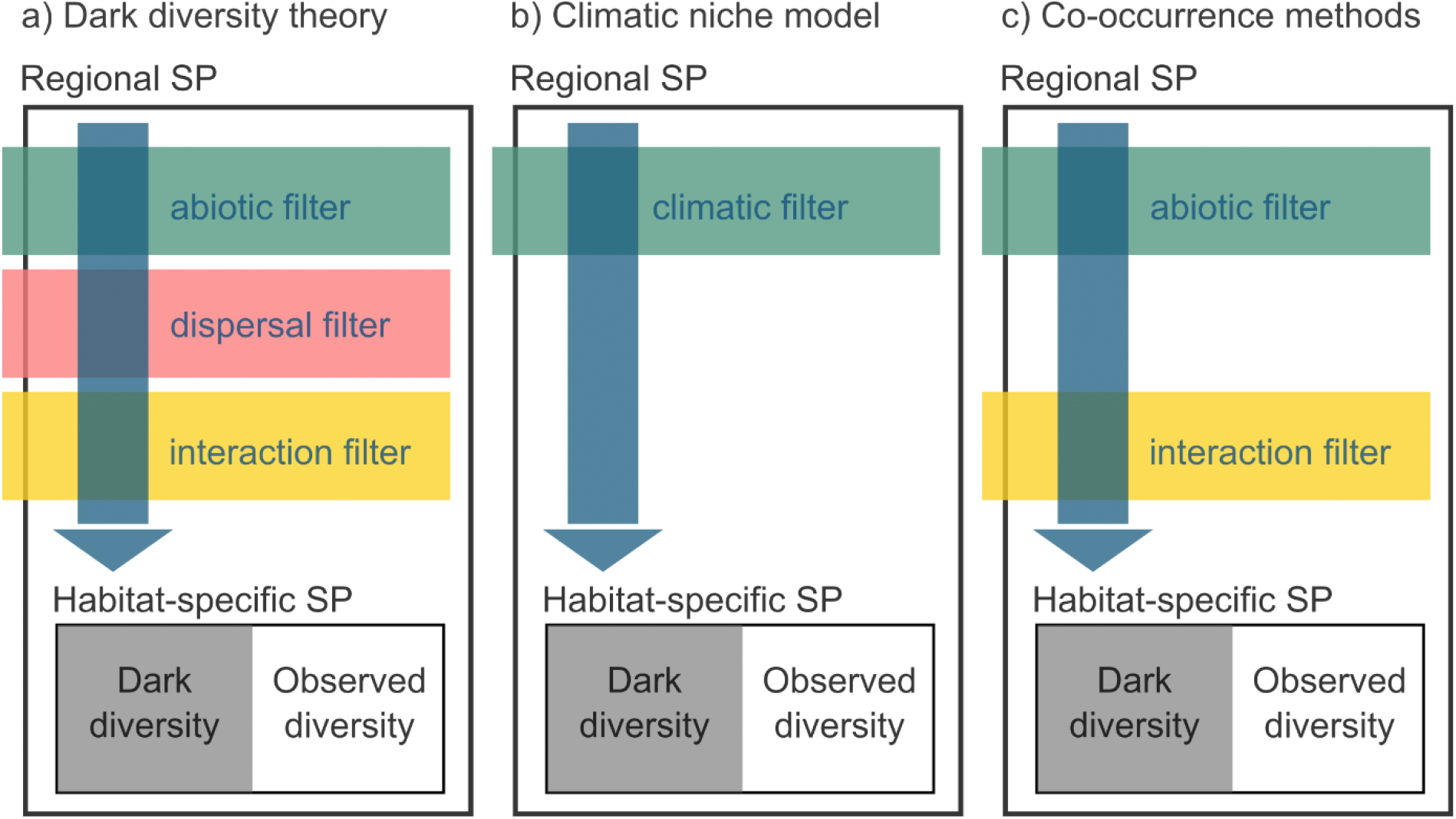
Schematic overview of three approaches to estimate the habitat specific species pool (SP). A) the theoretical concept of dark diversity, where the dark diversity is the non-observed set of species in a certain location, after filtering the regional species pool based on abiotic, dispersal and biotic interaction limitations. In b), dark diversity is calculated using climatic filtering of the regional species pool (e.g. using climatic niche models to estimate which species could occur at a certain location), while c) represents the commonly used co-occurrence based method, which integrates both abiotic and interaction filters. Figure adjusted from Stephenson (2016). The combination of dark- and observed diversity encompasses the habitat-specific species pool.

The dark diversity concept does not encompass the total regional species pool across different habitats but focuses on the environmentally filtered, or habitat-specific, regional species pool (Lewis et al., 2017). Combining this habitat-specific regional species pool with the local observed species composition can result in an estimate of the dark diversity (Figure 1). However, there are several methods that use different biotic and abiotic filters to estimate the habitat-specific species pool (Figure 1). Depending on the method, different outcomes will be obtained, as explained below. One of the main benefits of the dark diversity concept is that it allows us to compare the status of biodiversity in different habitats or ecosystems even if the local diversity differs by multiple orders of magnitude (Pärtel et al., 2011).

From a conservation perspective, the fact that species can be part of a community does not always mean that they should be, since not all species are desirable (Lewis et al., 2017). By only looking at suitable environmental conditions, non-native species adapted to those conditions will, for example, also be included. Very often, however, established non-native species have become part of the habitat-specific regional species pool and can, as a result, also be part of the dark diversity in the focal site. Dark diversity can therefore not only be used to investigate unwanted absences and ways to mitigate them, but also to prevent unwanted presences when non-native species are considered part of the habitat-specific regional species pool (Lewis et al., 2017). Monitoring both the dark and observed diversity can then give an early warning signal when non-native species are part of the dark diversity of a location, i.e., even before they are entering the observed diversity. Additionally, the approach can also inform us when species are leaving the dark and entering the observed diversity in response to, for instance, restoration actions (Lewis et al., 2017).

Estimating dark diversity is not straightforward but can be done in multiple ways (Lewis et al., 2016, Figure 1). The difficulty lies in estimating the habitat-specific species pool, which is, as explained above, the set of species in a region that can persist in the environmental conditions of the target site (Pärtel et al., 2011). It encompasses both the observed and dark diversity of a habitat. One could perform extensive sampling of habitat types in a region to estimate the habitat-specific species pool of each habitat type but this can be costly and time-consuming (de Bello et al., 2016). Therefore, computational approaches are often implemented. Most commonly, two types of methods are used to estimate the habitat-specific species pool, either (1) based on the abiotic niche of the species (e.g., using ecological indicator values or species distribution models) or (2) based on metrics of species’ co-occurrence with other species in the region (e.g., the Beals’ probability index or the hypergeometric method) (Lenoir et al., 2010; de Bello et al., 2016; Carmona and Pärtel, 2020).

Ecological indicator values are often used as an estimate for a species’ ecological requirements. The approach allows to identify species from the regional species pool along environmental gradients based on their ecological preferences (Ellenberg et al., 1991). A downside of this method is the difficulty of defining the realized niche of species since such indicator values are rough estimates of the niche optimum along a few specific ecological gradients, often based on expert knowledge (Lewis et al., 2016). Potentially more accurate approaches based on abiotic conditions make use of habitat suitability models to estimate species’ environmental niches (Guisan & Thuiller, 2005). These models can be used to determine the environmental conditions suitable for a species (Parolo et al., 2008). In this method, the accuracy of the models highly depends on the resolution as well as on the selected set of environmental data (de Bello et al., 2016).

In both the above-mentioned methods, the aim is to estimate the suitability of a location based only on the environmental niche of the species, regardless of the other species co-occurring in said location. By contrast, one could also estimate the potential of finding a species at a certain location based on the presence of its associated species. The Beals’ probability index can be used to calculate species co-occurrence patterns (Beals, 1984). It uses the idea that the presence of a species that is frequently found together with another species could indicate shared suitable abiotic conditions (Ewald, 2002). If the associated species of a target species are observed, but the target species itself is not, it is part of the dark diversity. The hypergeometric method works similarly by verifying if certain species associations occur more often than predicted by chance and by estimating the dark diversity of a given species at a location from the likelihood of its co-occurrence with species present at that location (Carmona & Pärtel, 2020). The advantage of these approaches is that one only requires species observations and no environmental conditions. However, the prediction of the probability of a given species to belong to the dark diversity is dependent on the distribution of other species, which is especially challenging for species that are not strongly confined to particular communities or for environments where traditional communities and thus species associations are truncated (e.g., due to habitat disturbances).

All these approaches have in common that they help identify species that are part of the habitat-specific species pool but that are not recorded in the observed diversity at a certain location, making them part of the dark diversity (Figure 1; Pärtel et al., 2011). In this paper, we applied both niche- and co-occurrence-based methods to estimate dark diversity in a case study on the subarctic tundra around Abisko, northern Sweden. We then further explored the drivers behind the spatial patterns of this dark diversity. The concept of dark diversity is still in its infancy and therefore only a handful of studies have explored why species are part of the dark diversity, none of which were to our knowledge conducted in a tundra climate (Belinchón et al., 2020; Moeslund et al., 2017; Riibak et al., 2015). In this study, we wanted to unravel the drivers behind (1) why some locations are missing relatively more suitable species than others, and (2) why certain vascular plants of the Scandinavian tundra are more often absent from suitable locations than others.

In light of the first research question, we expected locations with a higher relative dark diversity, hereafter referred to as plot-level dark diversity (i.e., a higher percentage of missing species from the habitat-specific species pool) to: (1) appear at lower elevations, as more intense competition will exclude a higher proportion of species (Jones & Gilbert, 2016); (2) be at the extreme ends of disturbance gradients, based on the intermediate disturbance hypothesis (Lembrechts et al., 2014; Rashid et al., 2021); (3) be at the extreme end of low pH and/or moisture gradients, since such conditions can be tolerated by a few species only (Gough et al., 2000; Vonlanthen et al., 2006); or (4) have low observed species richness, as these locations will be dominated by highly competitive species preventing specialist species from co-occurring (Pellissier et al., 2010). Of course, these factors would act in addition to the stochasticity that always explains part of the variation in species occurrences at small spatial scales (Mohd et al., 2016).

Secondly, we predict that plant species with a higher dark diversity probability, hereafter referred to as species-level dark diversity (i.e., absent in a higher percentage of plots where they were predicted to occur) to: (1) have a higher specific leaf area (SLA), since the Scandinavian tundra is a nutrient-poor environment (Westoby, 1998); (2) have a lower maximum vegetative plant height, as smaller plants would be more easily outcompeted in plots were they could theoretically occur; (3) have a higher seed mass or short-distance dispersal, since these are (loosely) correlated to a limited dispersal ability and lower seed abundance, which decreases the number of successful dispersal events (Howe & Smallwood, 1982; Ozinga et al., 2005); (4) be more recently introduced in the region, as non-native species have a more limited distribution and show possible time-lags in niche filling (Alexander et al., 2016; Crooks, 2005); or finally, (5) be associated with arbuscular mycorrhizal (AM) or ectomycorrhizal (EcM) fungi, as the native vegetation in the region is dominated by ericoid mycorrhizal (ErM) species (Clavel, 2022 unpubl.; Finlay, 2008; Tedersoo, 2017).

## 2 Materials and methods

### 2.1 Study area

The field data collection was performed in July and August 2021 in the Abisko area, northern Sweden (68°21’N, 18°49’E). The region has a subarctic montane climate with an average annual temperature of −0.6°C (1913-2020, although average annual temperatures have not dropped below 0°C since 2011) and average annual precipitation of 310 mm (Abisko Scientific Research Station, 400 m above sea level (a.s.l.); https://polar.se/). The soil is comprised of till, colluvium, and glacio-fluvial deposits (Callaghan et al., 2013). At high elevations, the area is covered in snow for about 27 weeks of the year (Callaghan et al., 2013). At low elevations, the vegetation is dominated by open birch forests (*Betula pubescens* Ehrh.), with additional presence of rowan (*Sorbus aucuparia* L.) and several willow species (*Salix* sp.). The understory vegetation often consists of heath species (e.g., dwarf birch (*Betula nana* L.), European blueberry (*Vaccinium myrtillus* L.) and black crowberry (*Empetrum nigrum* L.)), or meadow species (e.g., Alpine bistort (*Bistorta vivipara* L.), globeflower (*Trollius europaeus* L.) and Alpine saw-wort (*Saussurea alpina* DC.)) (Sonesson & Lundberg, 1974). Above the treeline (520 m a.s.l), the vegetation is dominated by alpine/arctic heathland species (e.g., blue heath (*Phyllodoce caerulea* L.), bog blueberry (*Vaccinium uliginosum* L.) and lingonberry (*Vaccinium vitis-ideae* L.)) (Kullman, 2015).

### 2.2 Field data collection

#### 2.2.1 Study sites

A total of 107 plots were surveyed in the vicinity of four mountain trails: Björkliden, Låktatjåkka, Nuolja, and Rallarvägen (Figure 2).

**Figure 2:**
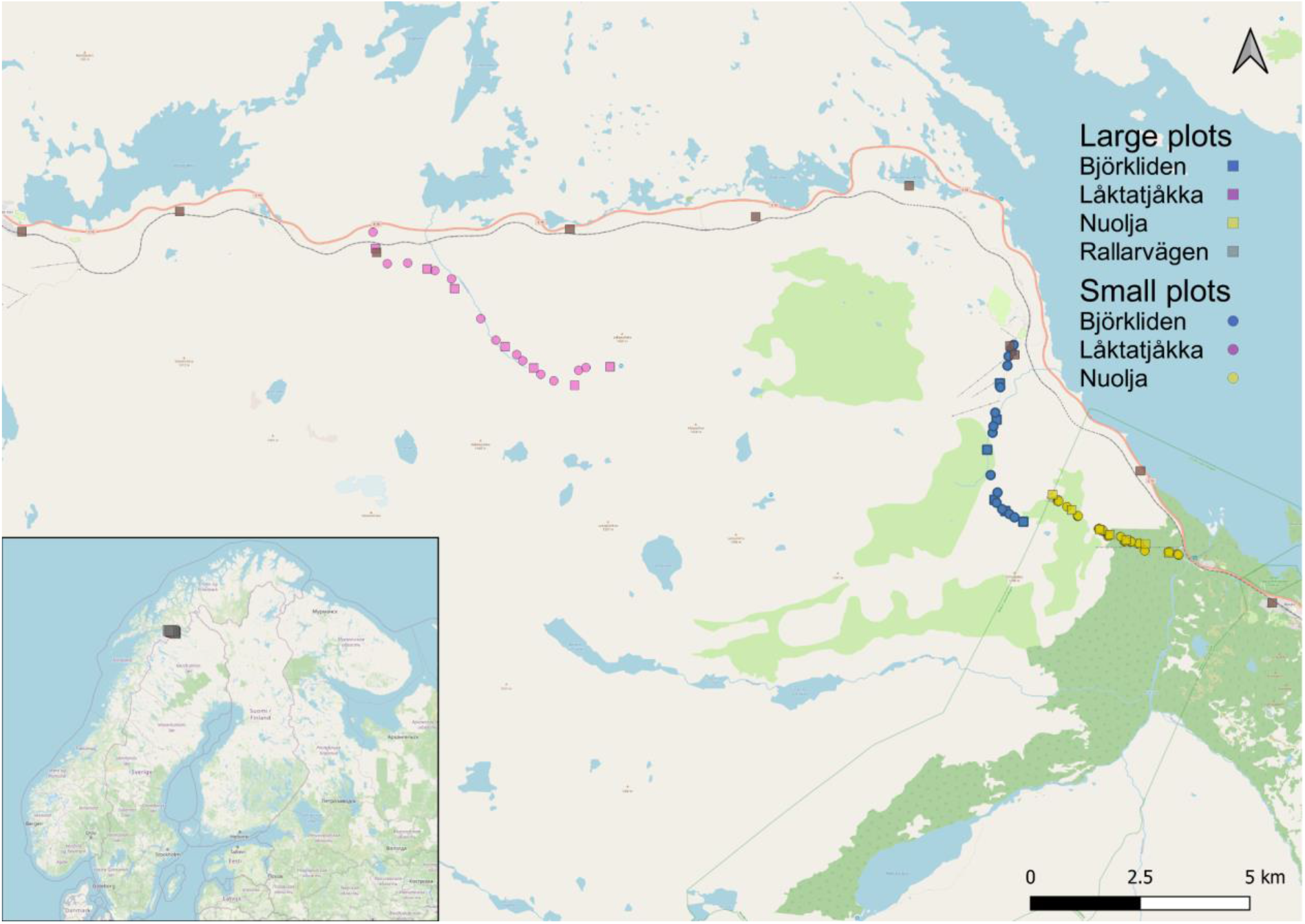
Map of the study area around Abisko, Sweden (grey dot on the inset), with 107 surveyed plots along the four hiking trails (colors) and the different survey methods (symbols).

Data from new and ongoing vegetation surveys were combined, with two different methodologies: 73 1 × 1 m^2^ plots from a long-term vegetation composition monitoring project in the area (hereafter called ‘small plots’), as well as 34 large (10 × 10 m^2^) plots established in the framework of the global DarkDivNet network (Pärtel et al., 2019). Of the 107 plots, two times 20 were located along trails close to Björkliden and around Låktatjåkka (Wedegärtner et al., 2022), 57 in the Abisko National Park on Mount Nuolja (MacDougall et al., 2021), and 10 along the Rallarvägen.

#### 2.2.2 Large plots

The vegetation monitoring method used in the large plots was based on the DarkDivNet protocol (Pärtel et al., 2019). The plots (10 × 10 m^2^) were placed at a 10 m perpendicular distance from the trail. In each plot, all vascular plants were recorded. Species were identified using the Fjällflora (Mossberg & Stenberg 2008). Observations that could not be identified to the species level (e.g., *Alchemilla* sp.) were removed from the species list and thus also from the regional species pool. Furthermore, following the DarkDivNet protocol, the maximum vegetative height (cm) was measured with a ruler for the tallest individual of each species in all plots.

In every plot, we visually estimated the cover (%) of total vegetation, bare ground, rock, litter, herbaceous vegetation, bryophytes, lichen, shrubs, and trees (> 200 cm). At the center of every plot, the exact location was recorded with a hand-held GPS receiver. Soil samples were collected using the protocol explained below (see 2.3).

#### 2.2.3 Small plots

The small plots were surveyed using the pin-point or point intercept method, which is often used to assess plant cover (Jonasson, 1988). A 1 × 1 m^2^ plot was placed at 10 m from the trail. In one plot, 100 pins were vertically dropped in 10-cm increments from left to right and top to bottom. With every pin-drop, we recorded the vascular plant species touching the pin, multiple recordings for the same species occurred when more than one individual of that species touched the pin. When the pin touched only the ground, the observation was categorized as either litter, bryophytes, bare soil, or lichen, a single hit was noted. Soil samples were collected using the same protocol as explained below (see 2.3).

### 2.3 Soil sample analysis

Soil samples were collected in 50 out of the 107 plots (both large and small plots). During sampling, the litter covering the soil was removed and a minimum of 300 g of soil was taken from the top 10 cm of the ground. Soil samples could not be collected along the Nuolja trail (57/107 plots) as this trail is in the Abisko National Park and no sampling permission was obtained in the year of the survey. However, 50 of these plots were long-term permanent plots for which soil pH measurements were available from previous soil sampling campaigns conducted in 2018 (using the same sampling and analysis procedure). The seven remaining plots were in very close (<10 m) proximity to small plots for which pH was measured in 2018, and we therefore used the mean pH of those plots. Ultimately, pH could be obtained for all but one plot, assuming that when largely undisturbed – as was the case in the system – pH-values would only change slightly over time.

All soil samples were stored in a fridge at 4°C until they were analyzed between September and December 2021 at the University of Antwerp, Belgium. To measure soil pH, 25 mL of a KCl solution was added to 10 g (9.9-10.1 g) of soil. The samples were put in a shaker for an hour and afterward rested for another 60 min. Then, soil pH was measured with a 914 pH/Conductometer by Metrohm© in the liquid layer at the top of the sample after shortly manually shaking the tubes.

### 2.4 Online data collection

#### 2.4.1 Gridded data products

To create the climatic niche models, we collected gridded climate data with a resolution of 30 arcseconds (c. 1 × 1 km^2^ at the equator) for the annual mean air temperature, annual precipitation, the mean maximum air temperature of the warmest month, and the mean minimum air temperature of the coldest month from CHELSA version 1.2, representing the long-term (1979-2013) climatic conditions (Karger et al., 2017).

Soil temperature estimates (i.e., annual mean soil temperature, mean soil minimum temperature of the coldest month and mean soil maximum temperature of the warmest month) were obtained from the SoilTemp global maps of soil temperature (Lembrechts et al., 2021). These maps calculated the soil bioclimatic variables using CHELSA monthly air temperature maps and the offset between gridded air temperature and in-situ soil temperature measurements from the SoilTemp database (Lembrechts et al. 2020). The gridded data, representative of the upper soil layer (top 5 cm), had the same resolution as the CHELSA data, namely 30”.

Elevation was extracted from the European Digital Elevation Model (DEM) with a resolution of 25 m, obtained from Copernicus Land Monitoring Service version 1.1 (European Union, 2021).

Lastly, the Topographic Wetness Index, a topographical proxy for soil moisture, was obtained from a TWI raster across Scandinavia, data acquired from Haesen et al. (2021). The TWI raster, which had a spatial resolution of 25 m, was generated using the method developed by Kopecký et al. (2021).

All data were extracted to the according coordinates of every plot in R version 4.2.1 (R Core Team, 2021) using the raster (Hijmans et al., 2012), sp (Pebesma et al., 2005), and GDAL (Mitchel et al., 2014) packages.

#### 2.4.2 Type of disturbance

For every plot, we assigned a type of disturbance based on its proximity to hiking trails, roads, and railroad. By visual assessment in QGIS, one of the three disturbance types (hiking trail, road or railroad) was assigned to every plot. All plots were close to hiking trails, yet whenever the railroad or a road was within 150 m of the plot, its impact was considered dominant, and the hiking trail classification thus overruled.

#### 2.4.3 Amount of bare ground

The amount of bare ground (%) as a proxy of disturbance was estimated or calculated for every plot. For the large plots, this was estimated from the percentage cover of litter and bare ground. This was calculated for the small plots by summing up all the pins that touched bare ground and litter, dividing this by the total number of pins in a plot.

#### 2.4.4 Plant functional traits

Average maximum vegetative plant height per species was calculated from the measurements done in the large plots.

The SLA for every species was retrieved from data collected in the framework of the Mountain Invasion Research Network (MIREN) in the region in 2017 (published as part of the Tundra Trait Team database (TTT); Bjorkman et al., 2018). The SLA was calculated as leaf area (cm^2^)/dry weight (g). Whenever there were two or fewer observations (30% of species, n=15) or no data at all (13%, n=6), the data was completed with values from the global Tundra Trait Team database (Bjorkman et al., 2018). Ultimately, for 28% (n=14) of the species, we extracted data from the TTT database and for 2% (n=1) of the species there was no data available. Within the TTT database, only single measurements on an individual - and thus not e.g., site-specific means - were used.

Average seed mass per species was obtained from the global TTT database or – if not available there - the LEDA Traitbase (Bjorkman et al., 2018; Kleyer et al., 2008). Again, from the TTT database, only ‘single measurements on an individual’ were used, whereas from LEDA only ‘actual measurements on individuals’ (no estimations) were used. The TTT database contained seed mass for 28% (n=14) of the species. An additional 47% (n=23) of the species could be supplemented with seed mass from the LEDA database.

The dispersal type per species was also retrieved from the LEDA Traitbase and used to categorize species according to their potential for long-distance dispersal (LDD) and short-distance dispersal (SDD) (Kleyer et al., 2008). For 86% (n=42) of the species this data was available and all were considered long-distance dispersers, hence this variable was not included in further analyses.

#### 2.4.5 Nativeness Index

We used a continuous rather than a binary measure of the status of a species within a region, to get a more accurate view of the history of the species. Our Nativeness Index (NI) used historical surveys from the Global Biodiversity Information Facility (<u>GBIF</u>) database. It considered the first year a species was observed (year first occurrence species) and the first year in which more than 50 species were observed in the region (year first survey). If the NI was close to 1, the species was already observed at the time of the first survey. As the value approached 0, the species was observed increasingly recently for the first time and was thus more likely to be non-native.

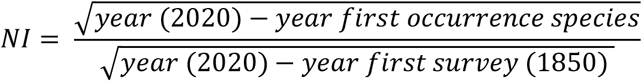

Square roots were used in the formula to give more weight to recent differences (e.g., a first observation in 2010 vs 2020 is considered a more substantial difference than one in 1900 vs 1910). The first occurrence and the year of the first survey were obtained using the *rgbif* package (Chamberlain et al., 2021).

#### 2.4.6 Mycorrhizal associations

The association of plant species with the main types of mycorrhizal fungi (AM = arbuscular mycorrhiza, EcM = ectomycorrhiza, ErM = ericoid mycorrhiza and NM = no mycorrhiza) was retrieved from the FungalRoot database (Soudzilovskaia et al., 2020). This data was available for 92% (n=42) of the species. For the remaining 8% (n=7) for which species-level information was absent, the mycorrhizal type was derived at the family level (i.e., the most common mycorrhizal type within the family).

### 2.5 Data-analysis

All statistical analyses were conducted in R version 4.2.1 (R Core Team, 2021).

#### 2.5.1 Dark diversity modeling

For further analysis, only the most common species, i.e., species with 10 or more observations, were included (n=49), as sufficient observations were needed to calibrate climatic niche models and build co-occurrence matrices. We then used the same dataset in four different approaches to estimate dark diversity.

##### Climatic niche modeling

The presence and absence of all species in every plot was used to make climatic niche models. For every species, a Generalized Linear Model was made, with a binomial distribution containing all climatic variables and their quadratic terms as explanatory variables (i.e., annual mean air temperature, annual precipitation, maximum air temperature of the warmest month, minimum air temperature of the coldest month, annual mean soil temperature, minimum soil temperature of the coldest month, and maximum soil temperature of the warmest month) and presence/absence (1/0) of a species per plot as the response variable. Multicollinearity was checked using the Variance Inflation Factor (VIF) from the *car* package (Fox & Weisberg, 2019) and variables that increased the VIF to 5 or more were removed. The final models contained: annual precipitation, minimum soil temperature of the coldest month, maximum soil temperature of the warmest month, and their quadratic terms. No further model selection was done as we were not interested in a model identifying the drivers of the species’ climatic niche, but rather wanted to approximate their climatic niche as consistently as possible.

To predict the probability of a species’ occurrence in a specific plot, the GLM was fit on the remaining plots (Lembrechts et al., 2019) and the probability was estimated for that specific plot. This leave-one-out procedure was then repeated for all plots and all species. We then calculated the relative dark diversity per plot by averaging the predicted presence of each absent species in a plot and the dark diversity probability per species by averaging the predicted presence of a species across all plots where it was absent.

The second method to estimate the dark diversity used the same climatic niche model as above. Yet, instead of continuous probability estimates, we converted niche model predictions into presence/absence estimates. For this, we calculated species-specific thresholds for presence using the function *ecospat.max.tss* from the *ecospat* package (Broennimann et al., 2022) which chooses the threshold that maximizes values for the True Skill Statistic (TSS), which assesses the accuracy of species distribution models (Allouche et al., 2006). Based on this, we created a binary dataset where the values below the threshold got a 0 (predicted to be absent) and the values above got a 1 (predicted to be present). Afterward, we removed the values where the species was observed to be present based on the vegetation surveys. To calculate the species-level dark diversity probability, we used the formula proposed by Moeslund et al. (2017), using the number of plot-level observations and predictions:

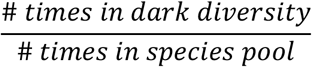

To calculate the relative plot-level dark diversity:

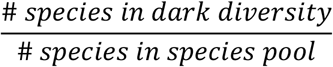

The habitat-specific species pool consisted of both the observed and dark species. Note that at the species level, we are estimating the probability that a species belongs to the dark diversity (dark diversity probability), while at the plot-level, we are estimating the percentage of species from the species pool that is absent (dark diversity *per se*).

##### Beals’ method

Two co-occurrence-based methods to estimate the dark diversity were used, with the first being the Beals’ index (Beals, 1984), as applied by Lewis, Szava-Kovats & Pärtel (2016). We first built a species co-occurrence matrix, then calculated the Beals’ index, using the *beals* function from the *vegan* package, for each species in every plot, excluding the focal species as suggested by Oksanen et al. (2022). The thresholds used to decide whether a species was part of the regional species pool were species-specific and defined as the 5th percentile of the Beals’ index value for the species (Gijbels, Adriaens & Honnay, 2012). Before calculating each threshold, the lowest value of the Beals’ index was determined among the plots containing occurrences of the species in question, and all plots with values below this lowest value were discarded (Moeslund et al., 2017). For each plot, the dark diversity then consisted of all species from the habitat-specific species pool, except those present (Pärtel, Szava-Kovats & Zobel, 2011). To calculate the plot- and species-level dark diversity probability the same formulae as for the species-specific threshold were used.

##### Hypergeometric method

The second method used to estimate the dark diversity was the hypergeometric method, as proposed by Carmona & Pärtel (2020). This method avoids the binary form in which dark diversity is often defined. The co-occurrence matrix used for the Beals’ method was also employed in this case. To get estimates of the dark diversity, we used the function *DarkDiv* from the *DarkDiv* package, with the argument ‘method’ containing ‘Hypergeometric’ (Carmona & Pärtel, 2020). We applied this method to all species in all plots for which we obtained a probability that the species could be present in that plot. Afterward, all values for plots where the species were observed to be present were removed. To calculate the relative plot-level dark diversity, per plot the mean was taken from the remaining values. The same was done for the species-level dark diversity, yet here the mean was taken per species.

#### 2.5.2 Drivers of relative plot-level dark diversity

To investigate why certain plots had a higher relative dark diversity, we created generalized linear mixed models (GLMMs) with a beta distribution and logit-link function using the *glmmTMB* package (Brooks et al., 2017), with predictions from each of the four dark diversity indices (the two approaches based on niche models, the Beals’ index, and the hypergeometric approach) as a response variable.

These plot-level models contained elevation, soil pH, type of disturbance, amount of bare ground, TWI, observed species richness and plot size as explanatory variables. The plots were situated along various trails. To account for this hierarchical sampling design, the model included a random intercept for plot number nested within trail identity. Multicollinearity and distribution of residuals were checked using the Variance Inflation Factor (VIF) and the *DHARMa* package (Hartig, 2022) and deemed not violated. Due to the low sample size, we limited ourselves to linear patterns and did not include two-way interactions since these more complex models could not converge. For the same reason, quadratic effects were not tested, even though theoretically they could be expected for pH and soil moisture. However, within our study system both the pH and moisture gradient only reached extreme values on one side of the gradient (e.g., highly acidic yet no highly basic soils).

No further model selection was performed (Hartig, 2018). The variance explained by the full model was obtained using the *performance* function from the *performance* package (Lüdecke et al., 2021). To determine the proportion of explained variance of every variable, we followed a variation partitioning approach. First, the variance of the full model was calculated. Afterward, for every explanatory variable, a model was made consisting of all variables except the focal variable. By extracting the marginal R^2^ of the individual models from the R^2^ of the full model, the variance of the focal variable was obtained (Legendre & Legendre, 1998).

#### 2.5.3 Drivers of species-level dark diversity probability

To investigate why certain species had a higher dark diversity probability, we created general linear models (GLMs) with a beta distribution and logit-link function using the *betareg* package (Cribari-Neto & Zeilis, 2010) with predictions from each of the used dark diversity indices (based on the niche models, the Beals’ index, and the hypergeometric approach) as a response variable.

First, full models were made separately for each dark diversity index that contained the nativeness index, maximum vegetative plant height, specific leaf area, dispersal type, seed mass, and mycorrhizal association as explanatory variables and species-level dark diversity as the response variable. Assumptions of multicollinearity and distribution of residuals were tested and not violated. Here as well, two-way interactions could not be tested and no further model selection was performed (Hartig, 2018). Afterward, pairwise comparisons were conducted on the categorical parameters using the *emmeans* package (Lenth, 2022).

## 3 Results

### 3.1 Plot-level dark diversity

Depending on the method, we could explain between 39% and 80% of the variance in plot-level dark diversity. In one case (climatic niche models), elevation was responsible for the largest share, while in the three other cases (species-specific threshold, Beals’ index and hypergeometric method) species richness was the most dominant factor (Figure 3). On average across all models, elevation explained 11%, species richness 10%, and plot size, type of disturbance, amount of bare ground, pH and TWI an additional 2%, 2%, 3%, 4% and 1%. Note that due to the nature of the variance partitioning calculations, variances do not necessarily add up to the total variance of the full model.

**Figure 3:**
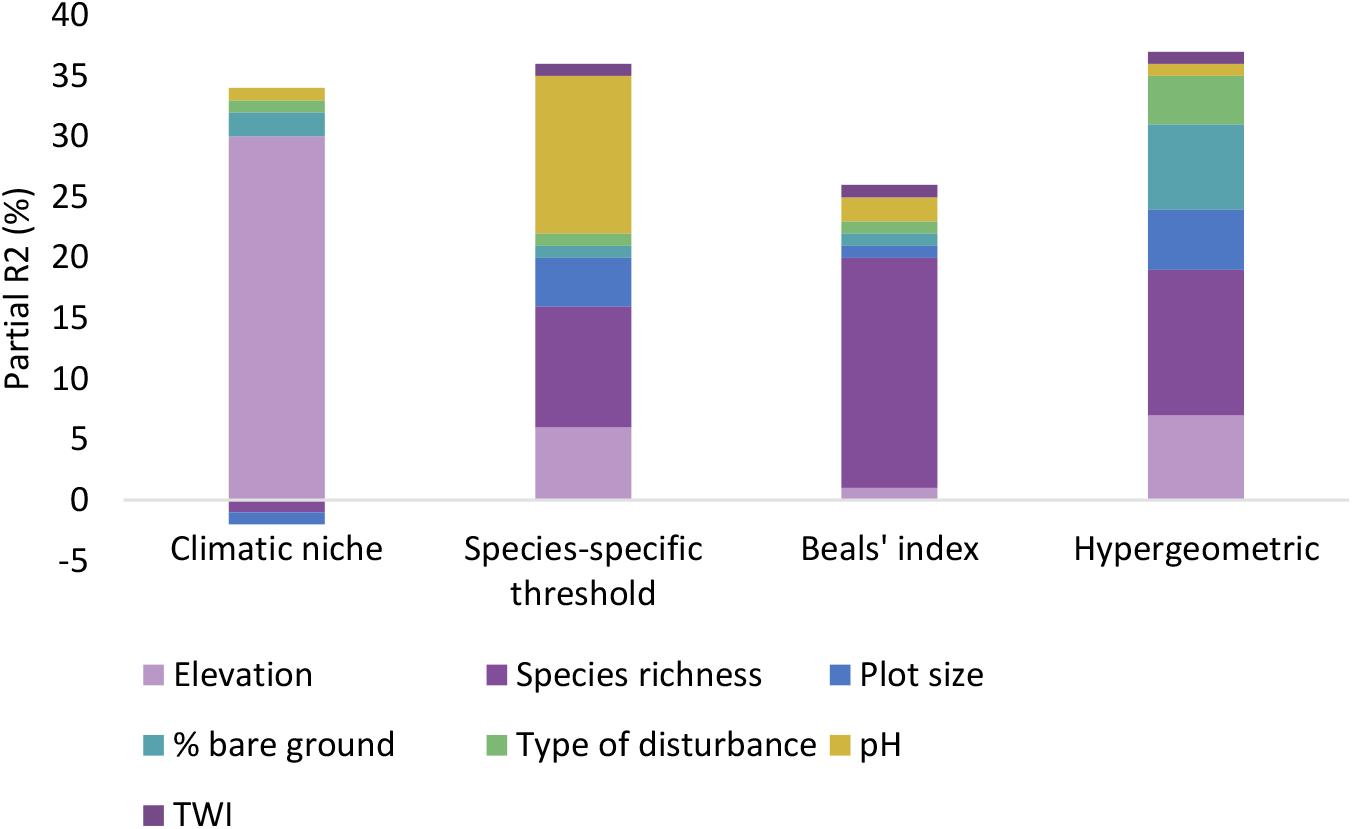
Variance partitioning (expressed in % and calculated using the marginal R^2^) of the different explanatory variables in the GLMMs of the plot-level analyses on the predictions of each of the four different dark diversity methods. TWI = Topographic Wetness Index.

In three out of the four methods used, the plot-level dark diversity decreased significantly across the elevation gradient (Table 1; Figure 4). Only in the model based on the Beals’ index did elevation not have a significant influence (Table 1; Figure 4D)

**Table 1:** Models explaining the plot-level dark diversity using the different dark diversity estimation methods: coefficients (p-values: * p≤0.05; ** p≤0.01; *** p≤0.001). The factor used for the intercept was allocated alphabetically and all other factors were compared to this baseline. CN = Climatic Niche models; SS = Species-specific threshold; Hyper = Hypergeometric method; Beals = Beals’ index; Elev = elevation; SR = Species Richness; TOD = Type Of Disturbance; TWI = Topographic Wetness Index.

| Model | Intercept<br>(Road) | Elev | SR | Plot<br>size<br>(10x10<br>m<sup>2</sup>) | TOD<br>Hiking<br>trail | TOD<br>Railroad | % bare<br>ground | pH | TWI | AIC |
| --- | --- | --- | --- | --- | --- | --- | --- | --- | --- | --- |
| CN | -0.327 | -<br>0.00<br>1*** | -<br>0.029**<br>* | 0.079 | 0.524 | -0.251 | -0.001* | -0.022 | -0.014 | -370 |
| SS | 3.00*** | -<br>0.00<br>1** | -<br>0.105**<br>* | -0.408* | 0.281 | -0.165 | -0.001 | -<br>0.262*<br>** | 0.038 | -140 |
| Hyper | 0.102 | -<br>0.00<br>1** | 0.001**<br>* | -0.183 | 0.425 | 0.001 | -<br>0.012**<br>* | 0.065 | -0.011 | -230 |
| Beals | 1.04* | 10<sup>-4</sup> | -<br>0.082**<br>* | -0.121 | -0.112 | -0.345 | 0.001 | 0.032 | 0.001 | -195 |

**Figure 4:**
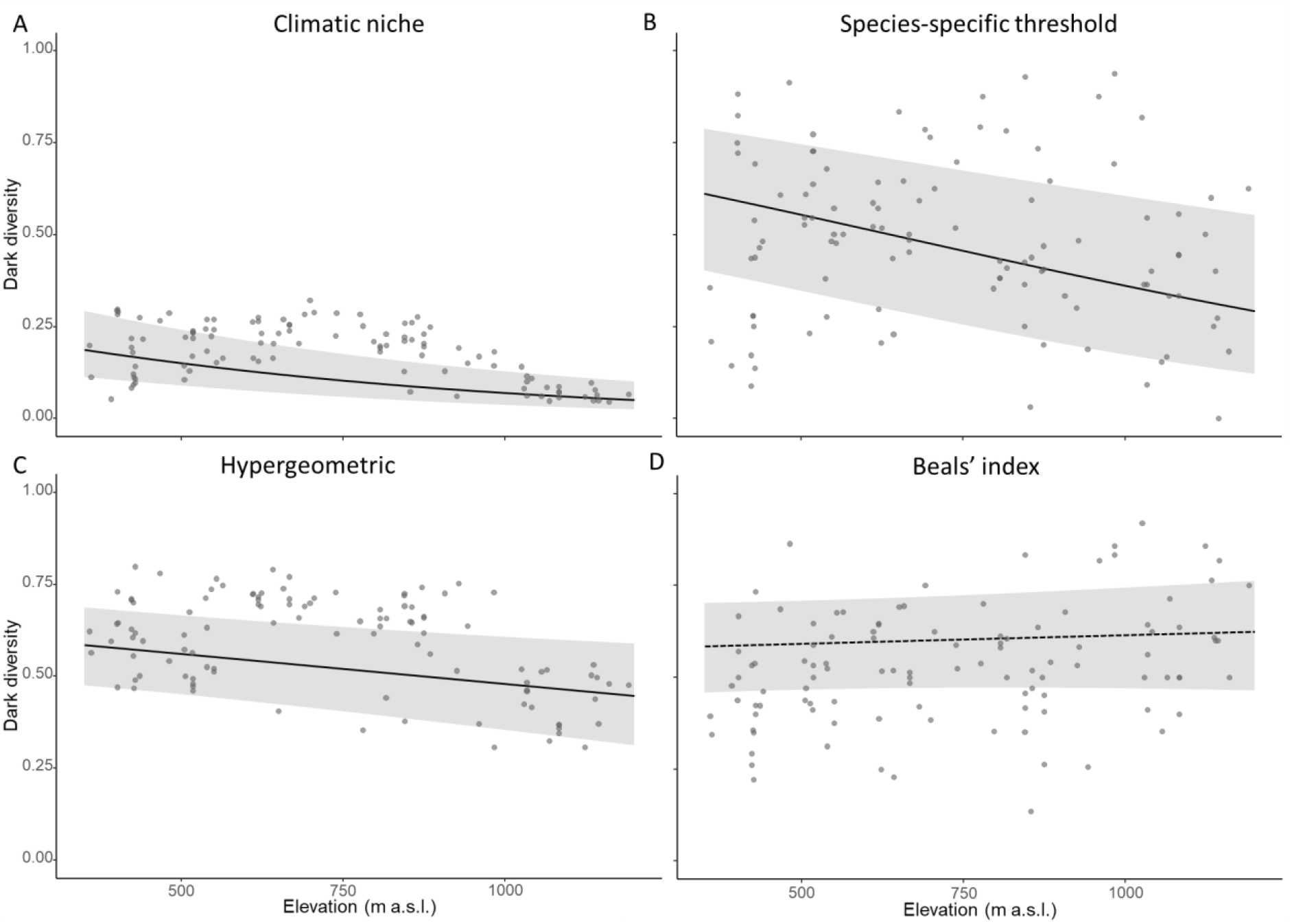
Marginal effects plots of the plot-level dark diversity as a function of elevation (m a.s.l.). The grey area indicates the 95% confidence interval, the grey dots are the raw data points, and the dotted line indicates non-significance. Dark diversity estimated using A) the climatic niche models, B) the climatic niche models followed by the species-specific threshold, C) the hypergeometric method and D) the Beals’ index.

The plot-level dark diversity decreased significantly with increasing species richness in three cases (Table 1; Figure 5A, 5B, 5D), yet increased significantly with increasing species richness when using the hypergeometric method (Table 1; Figure 5C).

**Figure 5:**
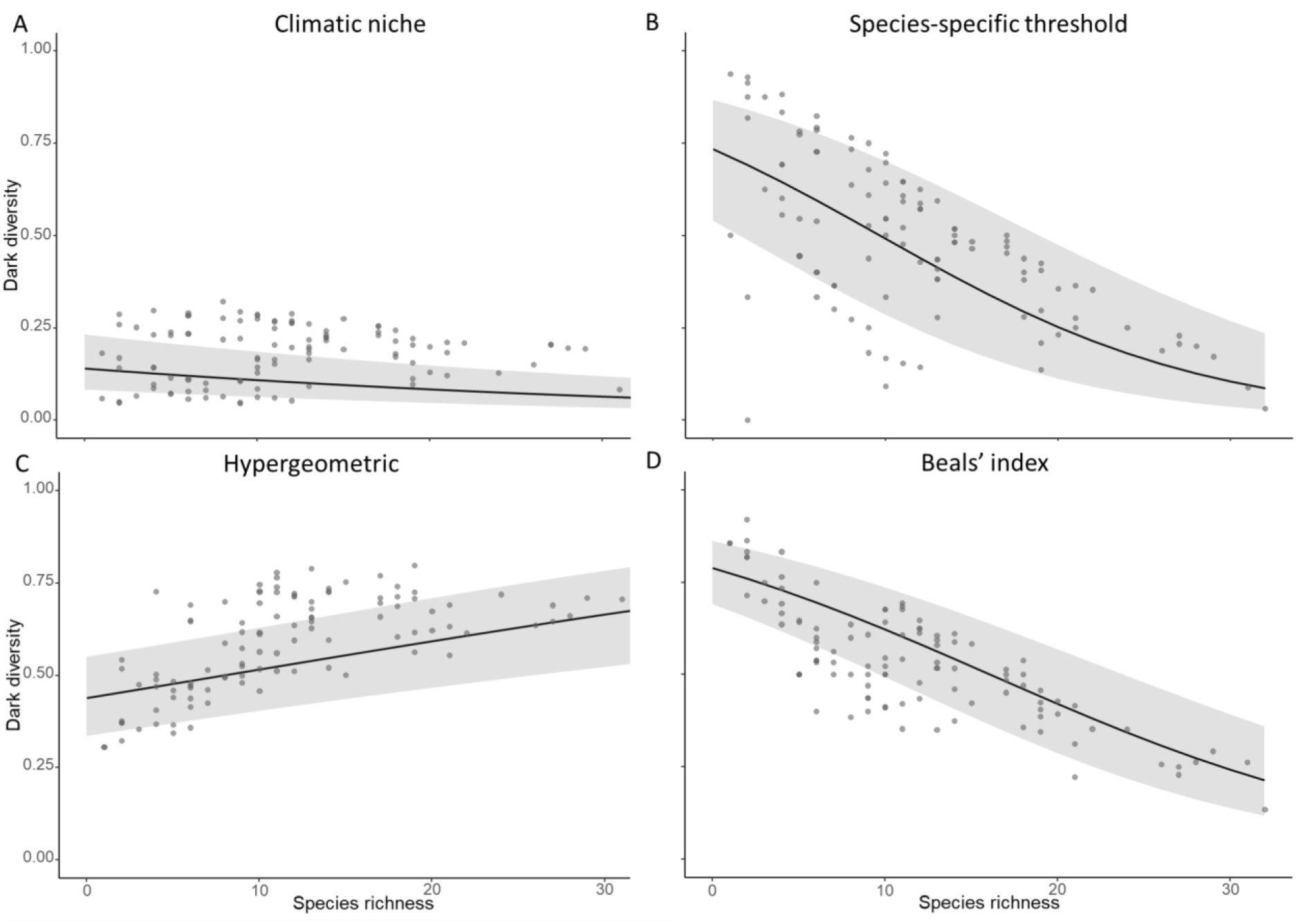
Marginal effects plots of the plot-level dark diversity as a function of species richness. The grey area indicates the 95% confidence interval, and the grey dots are the raw data points. A) the climatic niche models, B) the climatic niche models followed by the species-specific threshold, C) the hypergeometric method and D) the Beals’ index.

Moreover, the dark diversity decreased significantly with increasing amounts of bare ground when using the hypergeometric method (Table 1; Figure 6B) and the climatic niche models (Table 1; Figure 6A). In these models there was also some support that plots near a hiking trail showed a slightly higher dark diversity probability than plots closer to a road or railroad, although not significantly so (climatic niche: z = 1.65, *p = 0.099*; hypergeometric: z = 1.80, *p = 0.071*; Table 1; Figure 6C, D). The larger plots (10 × 10 m^2^) also had a significantly lower dark diversity compared to the smaller plots (1 × 1 m^2^) when using the climatic niche models followed by the species-specific threshold (Table 1; Figure 6E) and the hypergeometric method (Figure 6F), although in the latter not significantly (Table 1). Lastly, when using the climatic niche model followed by the species-specific threshold, the dark diversity decreased significantly with increasing pH (Table 1; Figure 6G), while the opposite was true for the hypergeometric method, albeit not significantly (z = 1.81; *p = 0.057*) (Table 1; Figure 6H).

**Figure 6:**
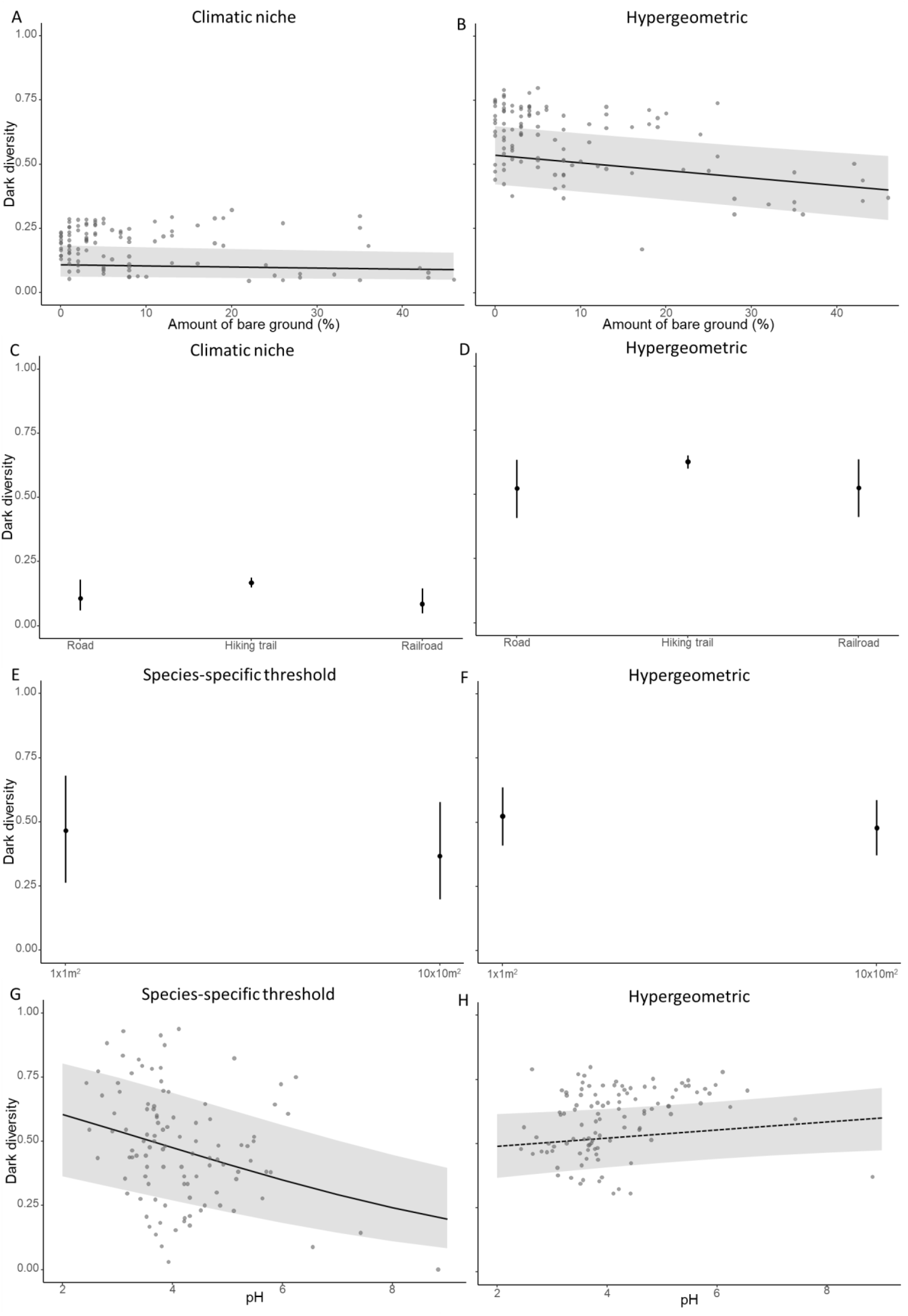
Marginal effects plots of the plot-level dark diversity as a function of the A/B) amount of bare ground, C/D) type of disturbance, E/F) plot size and G/H) pH. The black dots show the average dark diversity per individual factor whereas the error bars show the standard deviation. The grey area indicates the 95% confidence interval, the grey dots are the raw data points and the dotted line indicates non-significance. Dark diversity was estimated using A/C) the climatic niche models, E/G) the climatic niche models followed by the species-specific threshold and B/D/F/H) the hypergeometric method.

### 3.2 Species-level dark diversity

Depending on the method, we could explain between 15% and 45% of the variance in species-level dark diversity (Figure 7). In all cases, mycorrhizal association was responsible for the largest share (Figure 7). On average across all models, mycorrhizal association explained 18%, seed mass 8%, specific leaf area 3% and the nativeness index and the maximum vegetative plant height an additional 2%. Note that due to the nature of the variance partitioning calculations, variances do not necessarily add up to the total variance of the full model.

**Figure 7:**
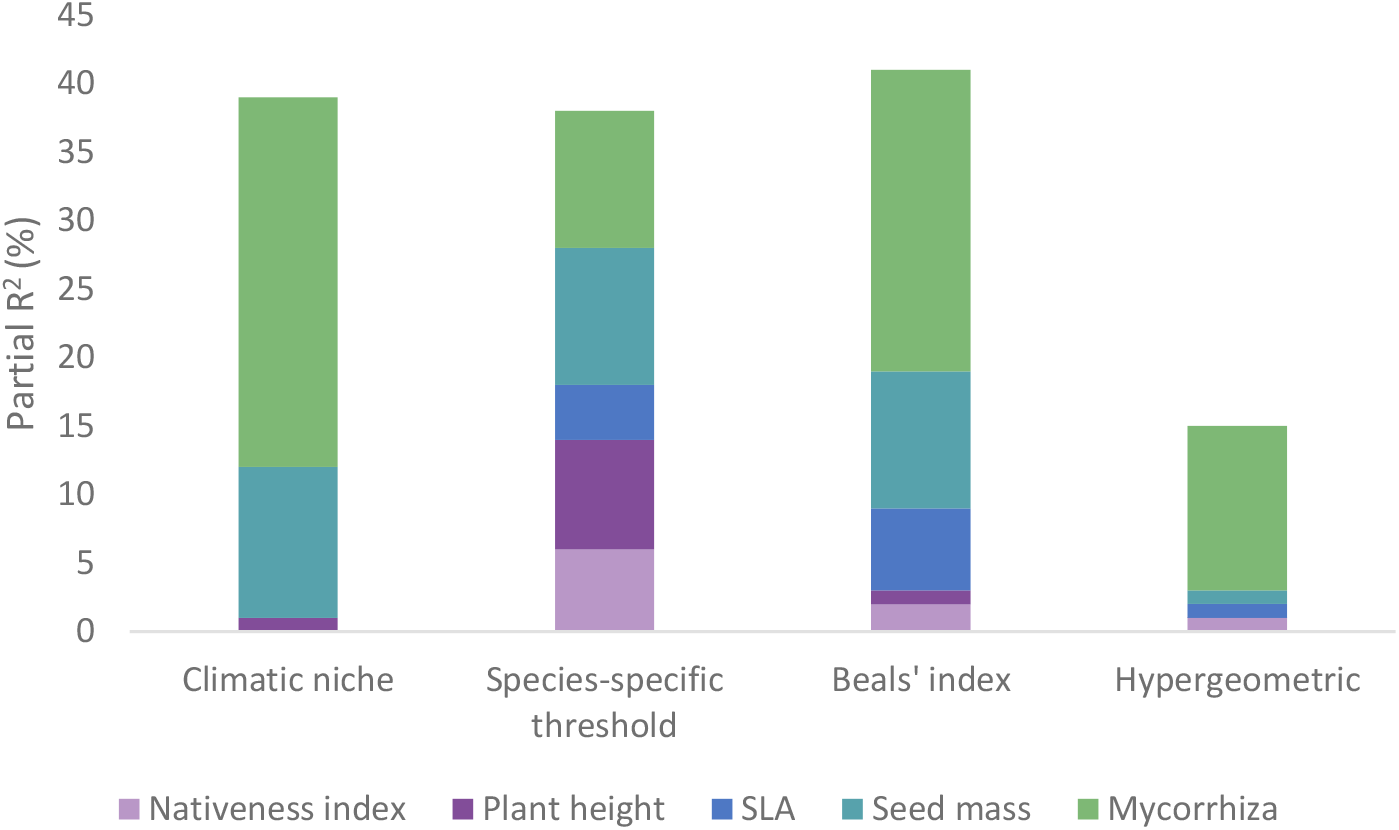
Variance partitioning (expressed in % and calculated using the marginal R^2^) of the different explanatory variables in the GLMMs of the species-level analyses on the predictions of each of the four different dark diversity methods. SLA = Specific Leaf Area.

Mycorrhizal status was the only significant parameter in the climate niche model approach, with ericoid mycorrhizae differing significantly from AM, EcM and NM (Figure 8; Table 2; Appendix S1). Species with a symbiotic ericoid mycorrhizal association had a significantly higher dark diversity than all other associations when using the climatic niche models (Table 2; Figure 8A). However, the opposite was true when using the three other methods (Table 2; Figure 8B, C, D). For the Beals’ index the contrast test also revealed ericoid mycorrhizae differing significantly from AM, EcM and NM (Figure 8D, Appendix S1). For the other two methods, the contrast test only showed a borderline significant difference between ErM and NM for the species-specific threshold and ErM AM for the hypergeometric method (Figure 8B, C; Appendix S1).

**Table 2:** Models explaining the plot-level dark diversity using the different dark diversity estimate methods: coefficients (p-values: * p≤0.05; ** p≤0.01; *** p≤0.001). The factor used for the intercept was allocated alphabetically and all other factors were compared to this baseline. CN = Climatic Niche model; SS = Species-specific threshold; Hyper = Hypergeometric method; Beals = Beals’ index; AM = Arbuscular Mycorrhiza; EcM = Ectomycorrhiza; ErM = Ericoid Mycorrhiza; NM = No Mycorrhiza; MVH = Maximum Vegetative plant Height; SLA = Specific Leaf Area; NI = Nativeness Index; SM = Seed Mass.

| Model | Intercept<br>(AM) | EcM | ErM | NM | MVH | SLA | NI | SM | AIC |
| --- | --- | --- | --- | --- | --- | --- | --- | --- | --- |
| CN | -0.043 | 0.108 | 1.17*** | -0.101 | $-10^{-4}$ | $-10^{-4}$ | -1.37 | -0.068 | -72 |
| SS | -5.96 | 0.453 | -0.916* | 0.208 | $-10^{-3}$ | $-10^{-3}$ | 6.95 | 0.082 | -11 |
| Hyper | 0.102 | -0.167 | -0.965* | -0.067 | $10^{-4}$ | 0.001 | 1.20 | -0.040 | -12 |
| Beals | 1.04* | 0.170 | -1.17*** | 0.193 | $-10^{-4}$ | -0.001 | -3.28 | -0.089 | -19 |

**Figure 8:**
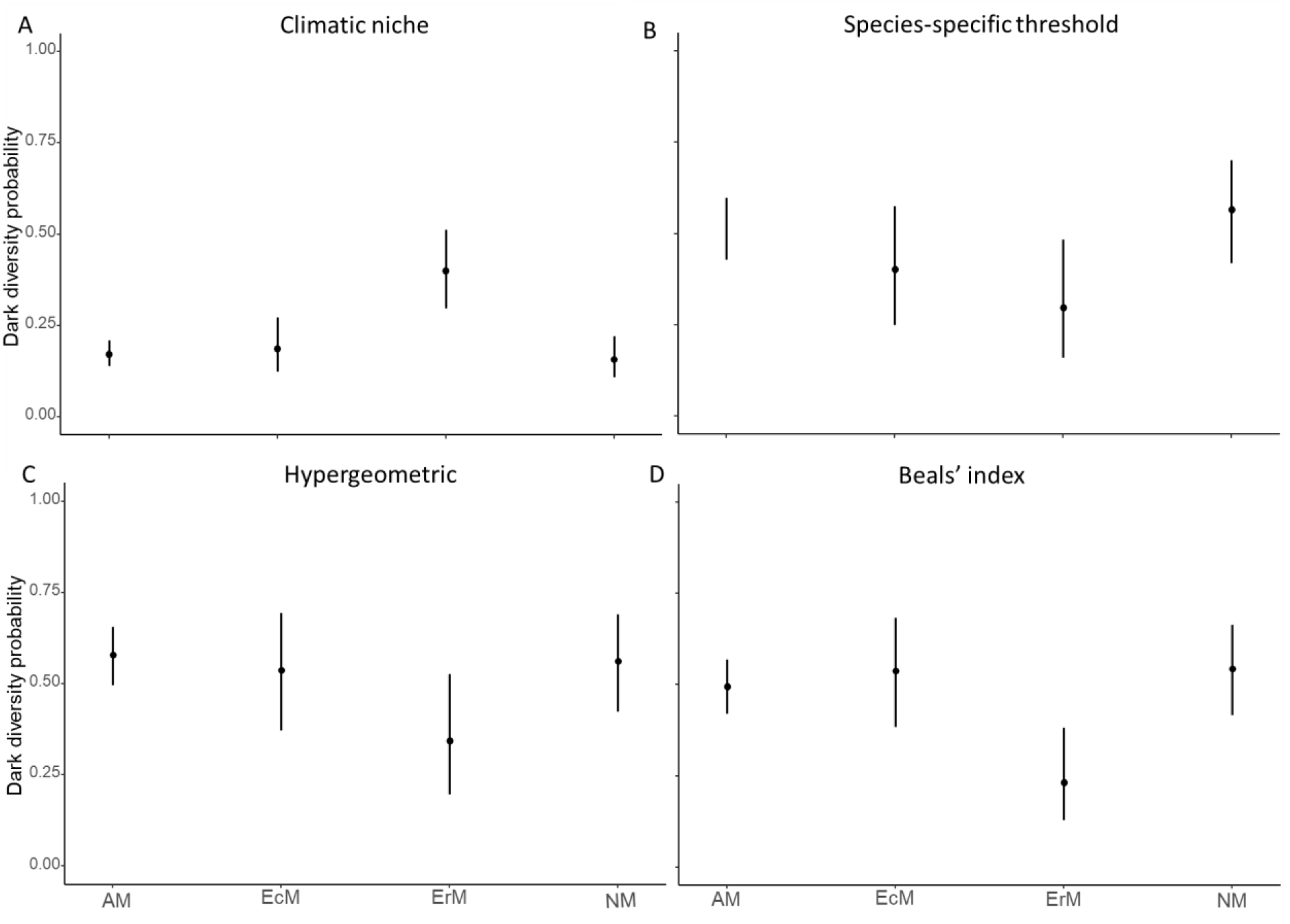
Prediction of the species-level dark diversity in relation to the type of mycorrhizal association based on the beta regression model. The black dots show the average dark diversity per individual factor whereas the error bars show the standard deviation. AM = Arbuscular Mycorrhiza; EcM = Ectomycorrhiza; ErM = Ericoid Mycorrhiza; NM = No Mycorrhiza. Dark diversity estimated using A) the climatic niche models, B) Species-specific threshold, C) the hypergeometric method and D) the Beals’ index.

Even though none of the other parameters had a significant influence on the species-level dark diversity, there is some support that the species-level dark diversity increased with increasing nativeness index (Figure 9B; z = 1.84, *p = 0.067*) and decreased with increasing maximum vegetative plant height (Figure 9A; z = −1.75, *p = 0.080*) when using the climatic niche model followed by the species-specific threshold, albeit both not significantly so. Lastly, there is also an indication (z = −1.86, *p = 0.062*) that the dark diversity decreased with increasing SLA when using the Beals’ index (Figure 9C).

**Figure 9:**
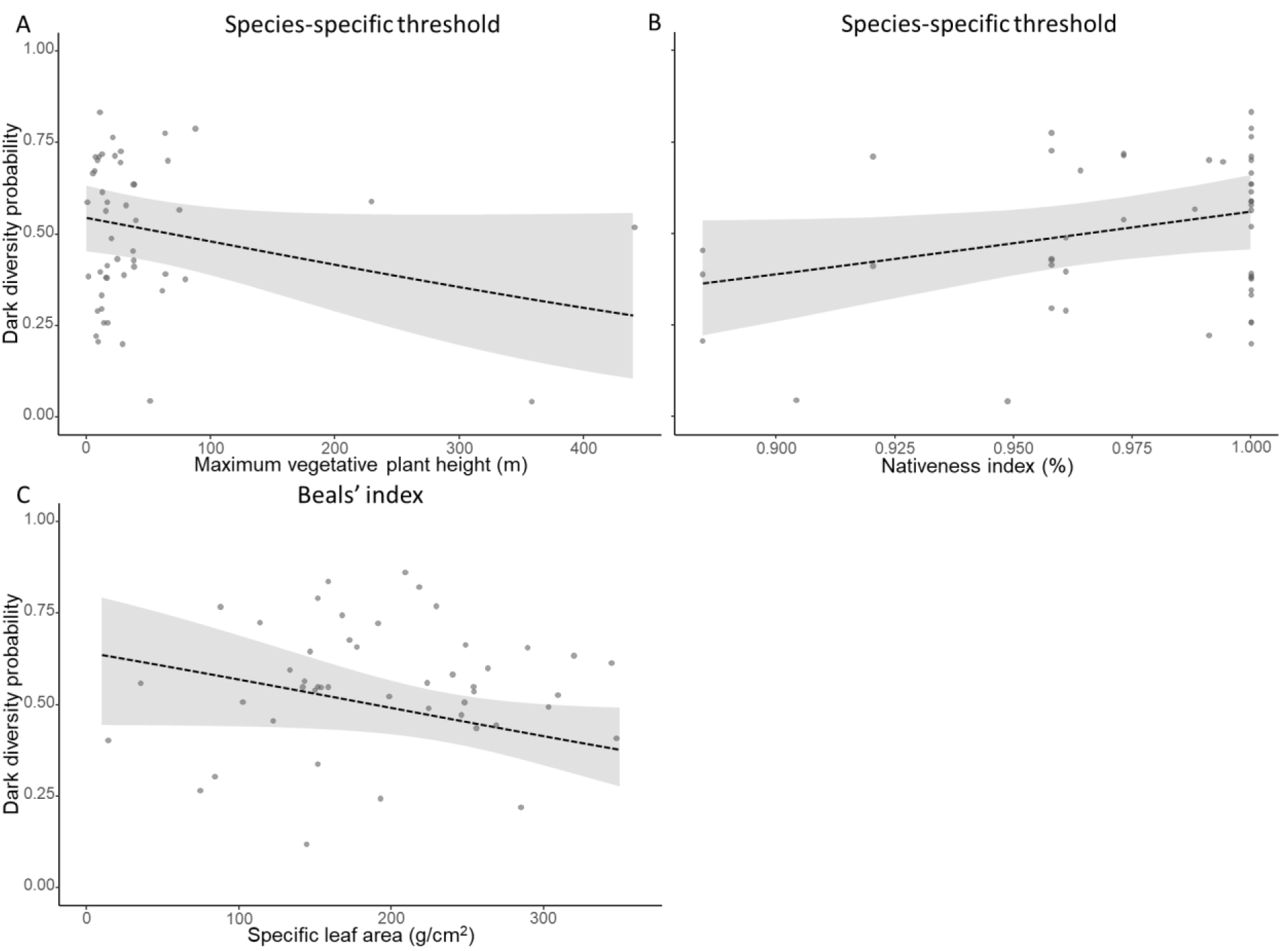
Marginal effects plots of the species-level dark diversity as a function of the A) Maximum vegetative plant height, B) Nativeness index and C) Specific leaf area. The dotted line indicates non-significance whereas the grey area indicates the 95% confidence interval and the grey dots are the raw data points. Dark diversity was estimated using A/B) the climatic niche models followed by the species-specific threshold, C) the Beals’ index.

## 4. Discussion

### 4.1. Plot-level dark diversity

We found relatively consistent patterns in the drivers of dark diversity at the plot-level, but much less consistency was observed at the species level. Plot-level dark diversity was most consistently related to elevation, with plots at higher elevations having a lower plot-level dark diversity - and thus fewer expected species missing - than plots at lower elevations. This was true for both niche-based methods as well as for the hypergeometric method, yet not for the Beals’ index, in which elevation was not significant. Such a decline with elevation is in line with ecological theory. Indeed, under harsh environmental conditions, competitive interactions are often replaced by mutualistic ones, or competition is at least lowered in intensity, thereby reducing the exclusion of less competitive species with a lower dark diversity as a result (Callaway et al., 2002; Klanderud, 2010; Lembrechts et al., 2018). Additionally, the presence of more ruderal and competitive species in the lowlands compared to the stress-tolerant species higher up in the mountains along roadsides also suggests that reduced competition can be one of the main drivers behind the lower dark diversity at higher elevations (Lembrechts et al., 2014). Furthermore, climatic conditions are usually milder in the lowlands, making them suitable for a broader set of species (Körner, 2021). Consequently, since more species can be present in these plots, it is also more likely that at least some of them are excluded, resulting in a higher number of species belonging to the dark diversity. As the co-occurrence-based metrics accounted for some of these factors (e.g., lower expectancy of species in plots dominated by species that traditionally outcompete them), it should come as no surprise that elevation was not significant in the model for the Beals’ index, and that the decline in dark diversity with elevation was the least steep for the hypergeometric method.

Species richness was identified as a key driver of plot-level dark diversity with all four methods. Its effect was negative for all but the hypergeometric method, thus largely following our hypothesis. In this system, plots with a low number of species are likely to be dominated by highly competitive species, which will prevent the establishment of several species that could in theory occur there (Pellissier et al., 2010). Indeed, plots with a low species richness in the study system were often dominated by black crowberry. It is an efficient competitor for nutrients, can grow on soils with low pH, and has allelopathic effects against seed germination and the growth of surrounding species (Tybirk et al., 2000), thus effectively referring several species from the regional species pool locally to the dark diversity. Nevertheless, it is possible that approaches based on species co-occurrences, such as the hypergeometric method and the Beals’ index, already account for this effect of competition. The latter potentially explains the observed positive relationship between species richness and dark diversity when using the hypergeometric method. Given the consistent negative relationship of dark diversity with elevation (see above), it is not unlikely that the effects of species richness and elevation would interact, with higher dark diversity especially in lowland plots with higher species richness, yet we lacked the sample size to explore such interactions statistically.

Finally, we found some support for the type of disturbance and percentage of bare ground as drivers of the plot-level dark diversity, while the results for soil pH were largely inconclusive. In two models, dark diversity tended to be lower in plots closer to the road - though none significantly so. Similarly, in two methods the dark diversity was found to decline significantly with increasing bare ground cover. Both point in the direction of a decreased dark diversity with increasing disturbance, a finding in line with other studies in the region. Indeed, bare ground cover could serve as a proxy for disturbance, with plots with virtually no bare ground usually being in late-successional stages (Lembrechts et al., 2014). In the Scandinavian tundra and other arctic, subarctic or alpine areas, late-successional stages – especially at low elevations - are often dominated by black crowberry, lingonberry or related species (Tybirk et al., 2000). As a result, these undisturbed plots are dominated by a few late-successional competitors, which can explain their higher dark diversity. More frequent disturbances, on the other hand, can result in more bare ground, which creates opportunities (empty niches) for species to establish and thus result in a lower dark diversity in these plots (Prieur-Richard & Lavorel, 2000). Lembrechts et al. (2014), for example, found in the same region a higher species diversity along disturbed roadsides than in natural communities owing largely to competitive release, a conclusion that found some support here in our observed lower dark diversity close to a road and the railroad, at least with some methods. Initially, we expected that disturbance by roads or railroads would result in a higher plot-level dark diversity because many stress-tolerant mountain species would not be able to cope with these disturbances (Auerbach et al., 1997). However, this would only be the case if the disturbance would be sufficiently intense to reduce species richness, which rarely – if ever – happened in our study plots, given our sampling focus on undisturbed vegetation only. Therefore, the same explanation as for bare ground stands, i.e., plots that were more disturbed created opportunities for species to establish, resulting in a lower plot-level dark diversity (Lembrechts et al., 2014).

### 4.2 Species-level dark diversity

Mycorrhizal association was the only variable with significant influence on the species-level dark diversity across all methods. However, while species with a symbiotic ericoid mycorrhizal association had a significantly higher dark diversity than all other associations when based on the climatic niche models, the opposite was true when using the other three methods. These contrasting results highlight the differences between these dark diversity estimation methods. In this system, the species with an ErM association (e.g. black crowberry and lingonberry) were virtually not climate-limited (occurring in 64 and 63 out of the 107 plots, respectively) and could in theory, based on their climatic niche, be present in all plots. Therefore, their dark diversity probability ended up being very high in any plot where they were absent, simply because of the underlying modelling approach. This issue was corrected by using species-specific thresholds, in which case mycorrhizal type was not withheld as significant.

These ErM-associated species not only dominated the landscape, but they were also often found in strong association with each other, resulting in clear predictions of their presence once one of them was present, when using the co-occurrence-based method. As their spatial connection in the field was so consistent, their estimated dark diversity using these methods ended up relatively low. Additionally, as ErM-fungi are the most dominant and widespread fungi in tundra regions (Tendersoo, 2017), in theory, there ought to be enough coverage of ErM-fungi so that the establishment of species associated with them should not be hampered. Consequently, there should be less reason for the species to be absent in areas where they could potentially occur than for AM-associated species (Tendersoo, 2017). All of this suggests that the observed higher dark diversity estimates for ErM-associated species by the climatic niche-approach are most likely a methodological artefact. These methodological issues could also explain why such little consistency was observed for the other studied drivers of species-level dark diversity, calling for caution when interpreting findings from any such dark diversity estimate separately.

### 4.3 Comparison of methods and uncertainties

In this paper, we estimated dark diversity using both niche-based and co-occurrence-based methods with both approaches often used interchangeably in literature. However, our results show that both approaches have significantly different assumptions and, as a result, get relatively incomparable results. Indeed, the niche-based approaches estimate the dark diversity as the set of species that could occur at a certain location based on their climatic niche or other environmental filters. The latter drivers are then often used as explanatory variables for the observed dark diversity, as done in the underlying study. For example, reduced competitive interactions in sites with larger percentages of bare ground would result in lower dark diversity, as is hinted at by our results.

Co-occurrence-based methods, on the other hand, estimate dark diversity simply by the neighbouring species with which a target species is usually associated. These approaches incorporate biotic interactions inherently in the dark diversity estimate. However, they do exclude species from the dark diversity for which the climatic conditions fall within their climatic limits, yet whose co-occurring species are also missing at a site. The latter could be especially problematic in diverse communities with high beta diversity, or areas with truncated, reduced, or novel communities as a result of anthropogenic land use or climatic changes (Christensen et al., 2021).

Perhaps more worryingly, within each type of dark diversity estimation method, results were not necessarily in agreement with each other. We found largely different findings, especially for species-level dark diversity, when using climatic niches with or without species-specific thresholds, as well as when using the hypergeometric method versus the Beals’ index. As such, our results highlight the need for caution when calculating and interpreting dark diversity estimates, as the conclusions depend heavily on the methodological decisions made, and methods should thus be tailored to the specific research questions.

Of course, several alternative methods could still be used to estimate dark diversity, and many adjustments to the methods used above could be proposed. For example, one could use global datasets such as GBIF to model the climatic niche, rather than data from the study region only. Using global datasets for such broader-scale niche models could result in a more accurate estimate of the climatic niche since the entire climatic niche could be modelled, rather than a truncated version as results from regional data. However, most of these global datasets lack absence data and presences are obtained using a wide variety of methodologies and spatial resolutions (Tessarolo et al., 2014), while abiotic data is at the global scale often only available at coarser resolution (Lembrechts et al., 2019). This could also make the predictions less accurate. Additionally, there is the possibility of mismatches, especially for rare species, since global datasets can be strongly spatially biased (Meyer et al., 2016). Therefore, predicting local climatic niches based on global data can make it more difficult to figure out whether the absences are due to a bias in the global dataset or the drivers under investigation.

The most promising avenue could perhaps come from an approach that combines both climatic niches with co-occurrences, such as joint Species Distribution Models (jSDMs; Pollock et al., 2014). This recent class of distribution models draws information from species co-occurrences and explains spatial variation in species distributions by extending standard species distribution models with species– species associations. Such an approach could potentially allow distinguishing through one model between absences driven by environmental unsuitability, biotic interactions, or other drivers. Nevertheless, Carmona & Pärtel (2020) did find that jSDMs could not outperform the hypergeometric method, yet they do substantially increase computational time.

### 4.5 Conclusions

The concept of dark diversity is still in its infancy, yet its contribution to understanding community completeness and nature conservation has already been shown to be significant (Lewis et al., 2017; Riibak et al., 2015). In this context, it is crucial to determine whether a species’ absence is a result of species-specific traits or plot characteristics, be it abiotic factors or biotic interactions, which is something traditional biodiversity studies that only focus on species presences cannot provide. We here compared different methodological approaches to estimate dark diversity and showed significant divergence in predicted drivers of dark diversity based on the method used, calling for caution when interpreting statistical findings on dark diversity estimates. Nevertheless, we can generally conclude that areas at low elevations, and, to a certain extent, with a low species richness or low levels of disturbance showed a higher plot-level dark diversity, largely due to natural processes such as competitive dominance.

## Acknowledgments

We thank the master students Renée Lejeune and Jasmine Spreeuwers for their assistance in gathering data during the summer of 2021. Additionally, we extend our appreciation to Stef Haesen for supplying us with the Topographical Wetness Index raster.

## Data availability statement

Data will be made available on Zenodo upon acceptance of the paper.

