## Supporting information Appendix S1 for "The drivers of dark diversity in the Scandinavian tundra are metric-dependent"

Appendix S1

*Table 1: Estimated marginal means and contrasts among the three mycorrhizal associations (p-values: * p≤0.05; ** p≤0.01; *** p≤0.001). P-values between 0.05-0.1 are shown between brackets. AM = Arbuscular Mycorrhiza; EcM = Ectomycorrhiza; ErM = Ericoid Mycorrhiza; NM = No Mycorrhiza*

| Model | Contrast | Estimate |
|---|---|---|
| Climatic niche | AM - EcM | -0.0158 |
| Climatic niche | AM - ErM | -0.229*** |
| Climatic niche | AM - NM | 0.0139 |
| Climatic niche | EcM - ErM | -0.214** |
| Climatic niche | EcM - NM | 0.0297 |
| Climatic niche | ErM - NM | 0.243*** |
| Species-specific threshold | AM - EcM | 0.112 |
| Species-specific threshold | AM - ErM | 0.217 |
| Species-specific threshold | AM - NM | -0.0517 |
| Species-specific threshold | EcM - ErM | 0.105 |
| Species-specific threshold | EcM - NM | -0.164[1] |
| Species-specific threshold | ErM - NM | -0.268 (*p = 0.086*) |
| Hypergeometric | AM - EcM | 0.0411 |
| Hypergeometric | AM - ErM | 0.235 (*p = 0.093*) |
| Hypergeometric | AM - NM | 0.0165 |
| Hypergeometric | EcM - ErM | 0.194 |
| Hypergeometric | EcM - NM | -0.0246 |
| Hypergeometric | ErM - NM | -0.219 |
| Beals' index | AM - EcM | -0.0424 |
| Beals' index | AM -ErM | 0.263* |
| Beals' index | AM NM | -0.0481 |
| Beals' index | EcM - ErM | 0.305** |
| Beals' index | EcM - NM | -0.00567 |

| | | |
|---|---|---|
| Beals’ index | ErM - NM | -0.311** |
